# Viperin inhibits cholesterol biosynthesis and interacts with enzymes in the cholesterol biosynthetic pathway

**DOI:** 10.1101/2021.02.19.431989

**Authors:** Timothy J. Grunkemeyer, Soumi Ghosh, Ayesha M. Patel, Keerthi Sajja, James Windak, Venkatesha Basrur, Youngsoo Kim, Alexey I. Nesvizhskii, Robert T. Kennedy, E. Neil G. Marsh

## Abstract

Many enveloped viruses bud from cholesterol-rich lipid rafts on the cell membrane. Depleting cellular cholesterol impedes this process and results in viral particles with reduced viability. Viperin (virus inhibitory protein endoplasmic reticulum-associated, interferon-induced) is an ER membrane-associated enzyme that when expressed in response to viral infections exerts broad-ranging antiviral effects, including inhibiting the budding of some enveloped viruses. Here we have investigated the effect of viperin expression on cholesterol biosynthesis. We found that viperin expression reduces cholesterol levels by 20 – 30 % in HEK293T cells. A proteomic screen of the viperin interactome identified several cholesterol biosynthetic enzymes among the top hits. The two most highly enriched proteins were lanosterol synthase and squalene monooxygenase, enzymes that catalyze key steps establishing the sterol carbon skeleton. Co-immunoprecipitation experiments established that viperin, lanosterol synthase and squalene monooxygenase form a complex at the ER membrane. Co-expression of viperin was found to significantly inhibit the specific activity of lanosterol synthase in HEK293T cell lysates. Co-expression of viperin had no effect on the specific activity of squalene monooxygenase, but reduced its expression levels in the cells by approximately 30 %. Despite these inhibitory effects, co-expression of either LS or SM failed to reverse the viperin-induced depletion of cellular cholesterol levels in HEK293T cells. Our results establish a clear link between the down-regulation of cholesterol biosynthesis and viperin, although at this point the effect cannot be unambiguously attributed interactions between viperin and a specific biosynthetic enzyme.

## Introduction

Cholesterol is a critical component of eukaryotic membranes and a precursor to many steroid hormones and bile acids (1). Cholesterol biosynthesis occurs on the cytosolic face of the endoplasmic reticulum (ER), primarily in the liver and intestines (2) and is one of the most extensively regulated biosynthetic pathways (3). The numerous regulation mechanisms include transcriptional up-regulation through the sterol regulatory element binding protein (SREBP) pathway (4), down-regulation through sterol- and isoprenoid-dependent degradation of 3-hydroxy-3-methylglutaryl CoA reductase (HMGR) (5, 6), transcriptional regulation of HMGR (7, 8), post-translational modifications of HMGR (9) and, more recently, direct sensing of cholesterol leading to proteasomal degradation of a key biosynthetic enzyme, squalene monooxygenase (SM), by mediated by the E3 ubiquitin ligase, MARCH6 (10–12). Cholesterol is also an important component of lipid rafts in cell membranes (1, 13), which have been implicated in a variety of diseases, including Alzheimer’s disease, prion diseases, and bacterial and viral infections (13). Notably, various enveloped viruses require cholesterol-rich lipid rafts to bud from the cell, thereby completing the viral replication cycle to produce infectious viruses (13–18).

Eukaryotic cells exhibit a wide variety of defenses against viral infections (19). As part of the innate immune response, the first line of defense against infection, production of interferons (IFNs) promotes the up-regulation of a wide range of genes to combat infection (20, 21). Viperin – Virus Inhibitory Protein, Endoplasmic Reticulum-associated, Interferon iNducible – (also known as cig5 and RSAD2) is strongly induced by type I IFNs (16, 22–24) and is associated with a wide range of antiviral properties (22). In humans, viperin is a ~42 kDa, 361 amino acid, ER-associated protein that, interestingly, is one of only eight radical-SAM enzymes identified in the human genome. The enzyme comprises three domains: an N-terminal amphipathic helix responsible for localizing the enzyme to the ER membrane, a conserved radical-SAM domain containing a canonical 4Fe-4S cluster binding motif (CxxxCxxC), and a C-terminal domain, largely responsible for binding the substrate (Figure 1) (23, 25, 26). Viperin catalyzes the dehydration of CTP to form 3’-deoxy-3’,4’-didehydro-CTP (ddhCTP) through a radical mechanism (27). This modified nucleotide has been shown to act as an effective chain-terminating inhibitor of some, but not all, viral RNA-dependent RNA polymerases (27) (Figure 1).

**Figure 1.**
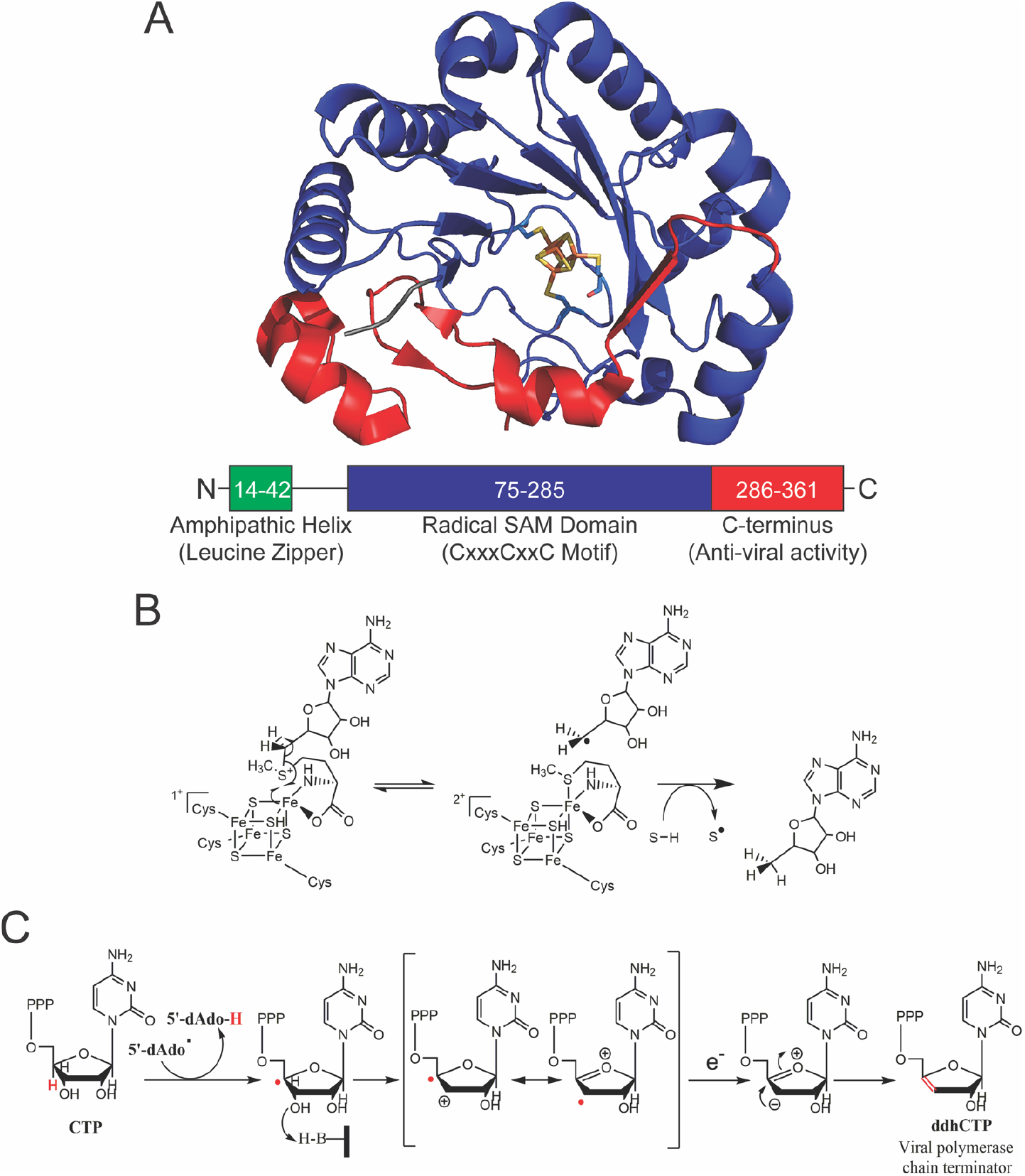
Structure and function of the viperin. **A.** Crystal structure of mouse viperin (PDB ID: 6Q2P) with the [Fe_4_S_4_] cluster bound, and its three domains highlighted. **B**. The general mechanistic scheme for the generation of 5’-deoxyadenosyl radical by radical SAM enzymes. **C.** Proposed mechanism for the formation of ddhCTP catalyzed by viperin.

Viperin is implicated in restricting a broad range of viruses including flaviviruses such as Dengue virus (DV) (28, 29), hepatitis C virus (HCV) (30–32), tick-borne encephalitis virus (TBEV) (33), West Nile virus (WNV) (29), and Zika virus (ZIKV) (33–35) and other types of viruses including human cytomegalovirus (HCMV) (22, 36, 37), human immunodeficiency virus (HIV) (38), and influenza A (18, 39). However, no common the mechanism has emerged by which viperin inhibits viral replication. Depending on the virus, the C-terminal domain has been reported to primarily facilitate viperin’s antiviral activity (21, 26, 28, 30, 40–42); in other cases, its antiviral effects have been attributed to the N-terminal helix or the radical-SAM domain (16, 18, 27, 30, 31, 41–49). The prevailing antiviral mechanisms proposed are localization to the ER and lipid droplets to inhibit viral budding (14, 16, 20, 31); inhibition of viral genome replication either by production of ddhCTP or interaction with viral replication complexes (27, 28, 30, 33, 40, 43, 44, 47, 50); stimulation of innate immune response pathways by interactions with intracellular signaling proteins (35, 49, 51, 52); and targeting of viral proteins for proteolytic degradation through the ubiquitination pathway (33, 43, 51, 53).

Viperin interacts with a diverse array of cellular and viral proteins. For example, viperin binds to the nonstructural proteins of various flaviviruses, leading to inhibition of genome replication and/or viral assembly (28, 30, 33, 40, 50). Additionally, viperin facilitates innate immune signaling in the Toll-like receptor-7/9 pathways through interactions with interleukin receptor-associated kinase 1 (IRAK1) and E3 ubiquitin ligase, TRAF6, to promote K63-linked polyubiquitination of IRAK1 by TRAF6, eventually leading to up-regulation of IFN expression(49, 51, 52). Although primarily localized to the ER and lipid droplets, viperin can be translocated to the mitochondria where it binds the mitochondrial trifunctional protein β-subunit (HADHB) and inhibits the thiolysis of β-ketoacyl-CoA esters thereby inhibiting fatty acid catabolism and decreasing cellular ATP levels (36, 37, 54).

Here we report studies that identify enzymes in the cholesterol biosynthetic pathway as novel targets for viperin. Our studies point to a new role for viperin in regulating cholesterol biosynthesis, which may explain the enzyme’s previously observed effect of inhibiting the budding of influenza A and other enveloped viruses from cells.

## Results

### Viperin expression reduces cholesterol biosynthesis

Various studies had previously linked viperin to the downregulation of cholesterol biosynthesis (39, 46, 55), but the effect of viperin expression of cellular cholesterol biosynthesis had not been investigated. To examine this question, HEK293T cells were grown in reduced serum media containing lipoprotein-depleted serum to minimize uptake of exogenous cholesterol (56). Cells were transfected with either viperin or an empty vector control and grown for a further 36 h before harvesting. Lipids were extracted from the cell pellets and total cellular cholesterol levels quantified by LC-MS, with the results normalized to total cellular protein. The cholesterol content of the control cells was 79.6 ± 2.1 nmol/mg of cellular protein (n = 4) whereas for cells overexpressing viperin was the cholesterol content 63.1 ± 3.6 nmol/mg (n = 4) (Figure 3). This represents a 21 % decrease in cholesterol which is both statistically significant, p = 0.008, and potentially biologically significant given that the statin-induced reduction of cholesterol biosynthesis is reported to substantially impair the replication of enveloped viruses such as influenza A and respiratory syncytial virus (57, 58). Cells cultured in media containing serum that was not lipoprotein-depleted exhibited no significant change in cellular cholesterol levels in response viperin expression, implying that viperin alters cholesterol biosynthesis rather than cholesterol uptake.

### The interactome of viperin includes several cholesterol biosynthetic enzymes

Viperin and many of the enzymes involved in cholesterol biosynthesis are associated with the ER membrane (16, 59), leading us to consider whether viperin may down-regulate cholesterol levels by inhibiting one or more of these ER-bound enzymes. To examine which, if any, of the cholesterol biosynthetic enzymes may interact with viperin we undertook a proteomic analysis to map the interactome of viperin and in the process obtain a more comprehensive inventory of the cellular proteins that interact with viperin. To accomplish this, a viperin construct bearing a N-terminal 3x-FLAG tag was transiently expressed in HEK293T cells, as described in the methods section. 42 hours post-transfection, the expressed viperin was immunoprecipitated using anti-FLAG antibody-conjugated magnetic beads. The immunoprecipitated proteins were subjected to on-bead reduction, alkylation, and tryptic-digestion followed by mass spectrometry of the alkylated tryptic peptides. An empty 3x-FLAG-tagged pcDNA3.1(+) construct was transfected in HEK293T cells to serve as a control. The MS analyses were performed on three biological replicates for both the viperin and empty vector control samples.

For the initial list of identified proteins, containing 3670 hits, the fold change in protein abundance in the viperin samples over the empty vector control was calculated, with protein abundance estimated using spectral counts (i.e., the total number of peptide-to-spectrum matches, PSM, per protein). The shortened list was then analyzed using the Significance Analysis of INTeractome (SAINT) software (60) via the Contaminant Repository for Affinity Purification (CRAPome; www.crapome.org) web resource (61) to exclude common, non-specifically bound proteins. For each identified protein in the dataset SAINT computes, via statistical modeling of spectral counts across bait purifications and the controls, a confidence score (probability) that it is an interactor of the bait protein. By setting the SAINT probability cut off to be ≥ 0.9, we obtained a list of 100 possible hits (Table S1). It was apparent from this initial list that proteins involved in steroid biosynthesis and lipid metabolism and were heavily represented.

Therefore, we next performed a pathway enrichment analysis on these hits using DAVID, v 6.8 (The Database for Annotation, Visualization and Integrated Discovery) software. Functional annotation of these proteins through KEGG pathway analysis showed the highest enrichment scores, 3.59 in DAVID at high stringency, were for enzymes in sterol biosynthesis pathways (Table 1, Figure 2). This observation provided initial support for the hypothesis that viperin inhibits one or more of the enzymes in cholesterol biosynthesis.

**Figure 2.**
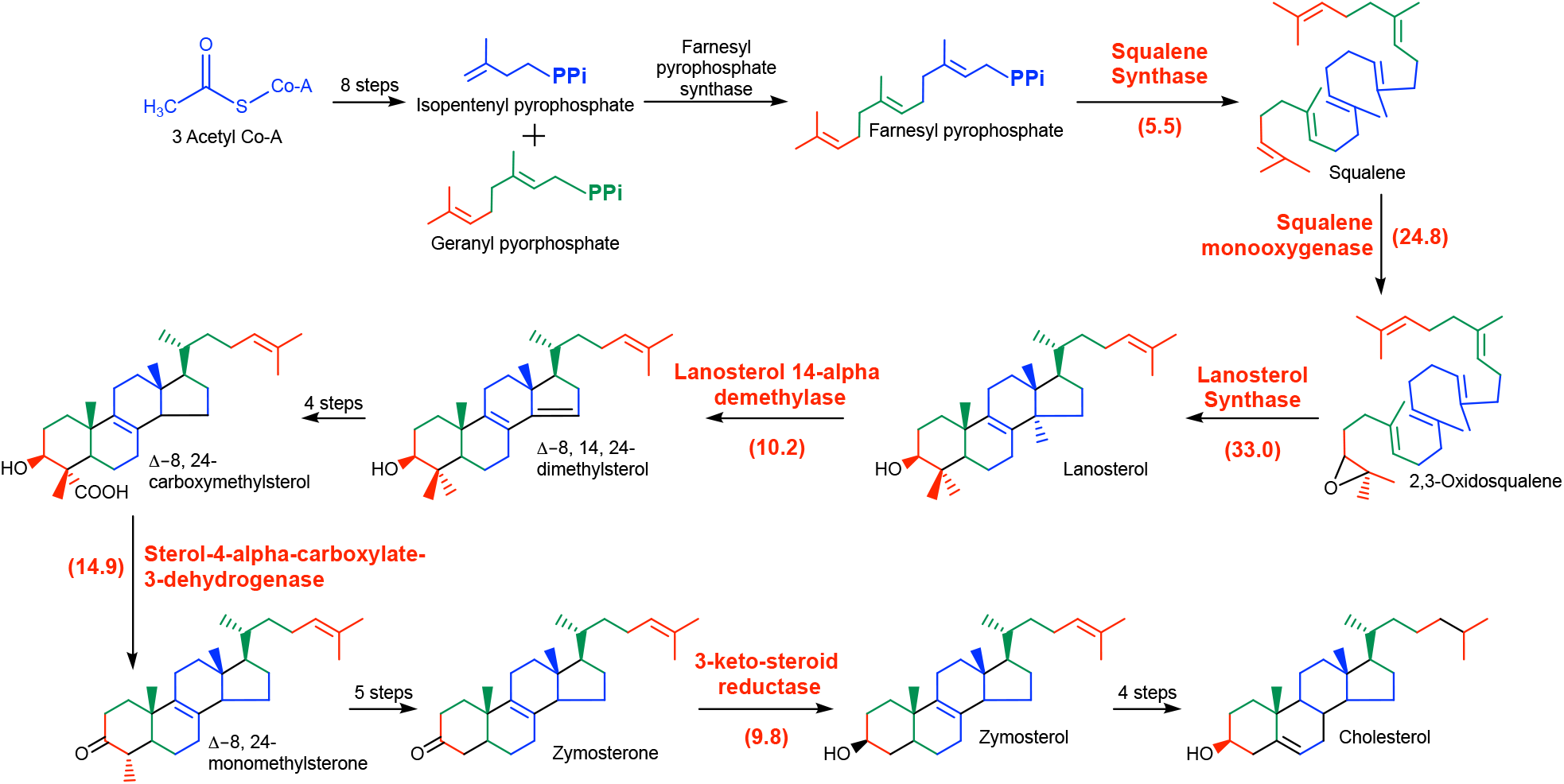
Enzymes from the cholesterol biosynthetic pathway identified in the interactome of viperin. An overview of the pathway is presented with enzymes identified as interacting with viperin highlighted in red, with enrichment factors given in parentheses.

**Figure 3.**
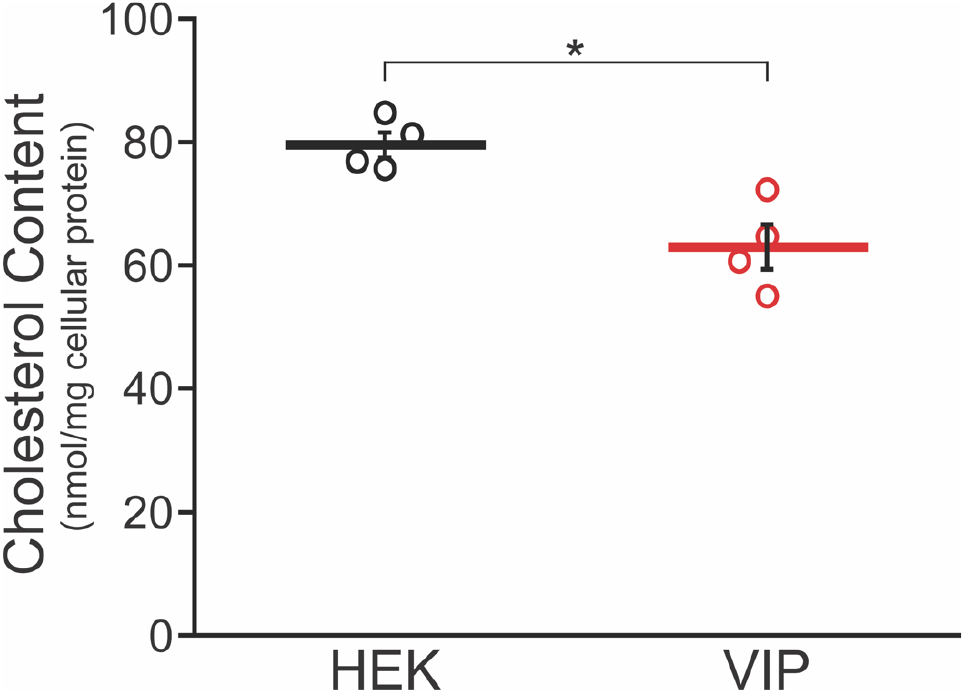
HEK293T cells expressing viperin exhibit reduced cholesterol levels. HEK293T cells were grown in reduced serum media containing lipoprotein-depleted serum and transfected with empty vector (HEK) or viperin (VIP) as indicated. Cholesterol was extracted from cell pellets and total cholesterol levels were quantitated via LC-MS using a standard curve generated using pure cholesterol. Results were normalized against total cellular protein content. *Denotes p value < 0.01.

**Table 1.** Cholesterol biosynthetic enzymes identified by the proteomic analysis of proteins that were co-immunoprecipitated by viperin. The fold enrichment represents the factor by which each protein was enriched in the sample with respect to the empty vector control.

| UniProtKB | Gene Name | Enzyme | Enrichment (fold) | SAINT Score |
| --- | --- | --- | --- | --- |
| P48449 | LSS | lanosterol synthase | 33.0 | 1 |
| Q14534 | SQLE | squalene monooxygenase | 24.8 | 1 |
| Q15738 | NSDHL | sterol-4-alpha-carboxylate 3-dehydrogenase, decarboxylating | 14.9 | 1 |
| Q16850 | CYP51A1 | lanosterol 14-alpha demethylase | 10.2 | 1 |
| P56937 | HSD17B7 | 3-ketosteroid reductase | 9.8 | 1 |
| P37268 | FDFT1 | squalene synthase | 5.5 | 0.97 |

### Viperin interacts with SM and LS to form a ternary complex

Many of the cholesterol biosynthetic enzymes have been shown to interact with each other in a functional complex (59) so it is likely that some of the enzymes identified by the screen are enriched through indirect interactions, rather than binding directly to viperin. Therefore, we focused on the two most highly enriched enzymes: squalene monooxygenase (SM) and lanosterol synthase (LS). Notably, SM and LS were both substantially enriched over other cholesterol biosynthetic enzymes and were the two most highly enriched proteins identified in the proteomic screen (Table S1). SM catalyzes the conversion of squalene to 2,3-oxidosqualene and LS catalyzes the subsequent cyclization of 2,3-oxidosqualene to lanosterol (Figure 2), which are key steps in sterol biosynthesis (2).

To validate the interactions between LS and SM with viperin, we transiently expressed these proteins in HEK293T cells and subjected them to co-immunoprecipitation analysis. Viperin, SM, and LS were transiently expressed with N-terminal 3x-FLAG, C-terminal V5, and C-terminal Myc epitope tags, respectively. Furthermore, because the N-terminus of viperin has proven important for its interactions with various other proteins (16, 30, 43, 47), we also co-expressed a truncated viperin construct lacking the N-terminal 50 residues (viperin-ΔN50) that localize viperin to the ER membrane to examine whether ER-localization was important in this instance. Co-immunoprecipitation experiments were performed using either viperin or viperin-ΔN50 as bait proteins with SM and/or LS serving as the prey proteins. Both viperin and viperin-ΔN50 were found to co-precipitate SM and LS when co-expressed individually. Consistent with this observation, viperin and viperin-ΔN50 also co-precipitated both SM and LS when all three enzymes were co-expressed (Figures 4A and 4B).

**Figure 4.**
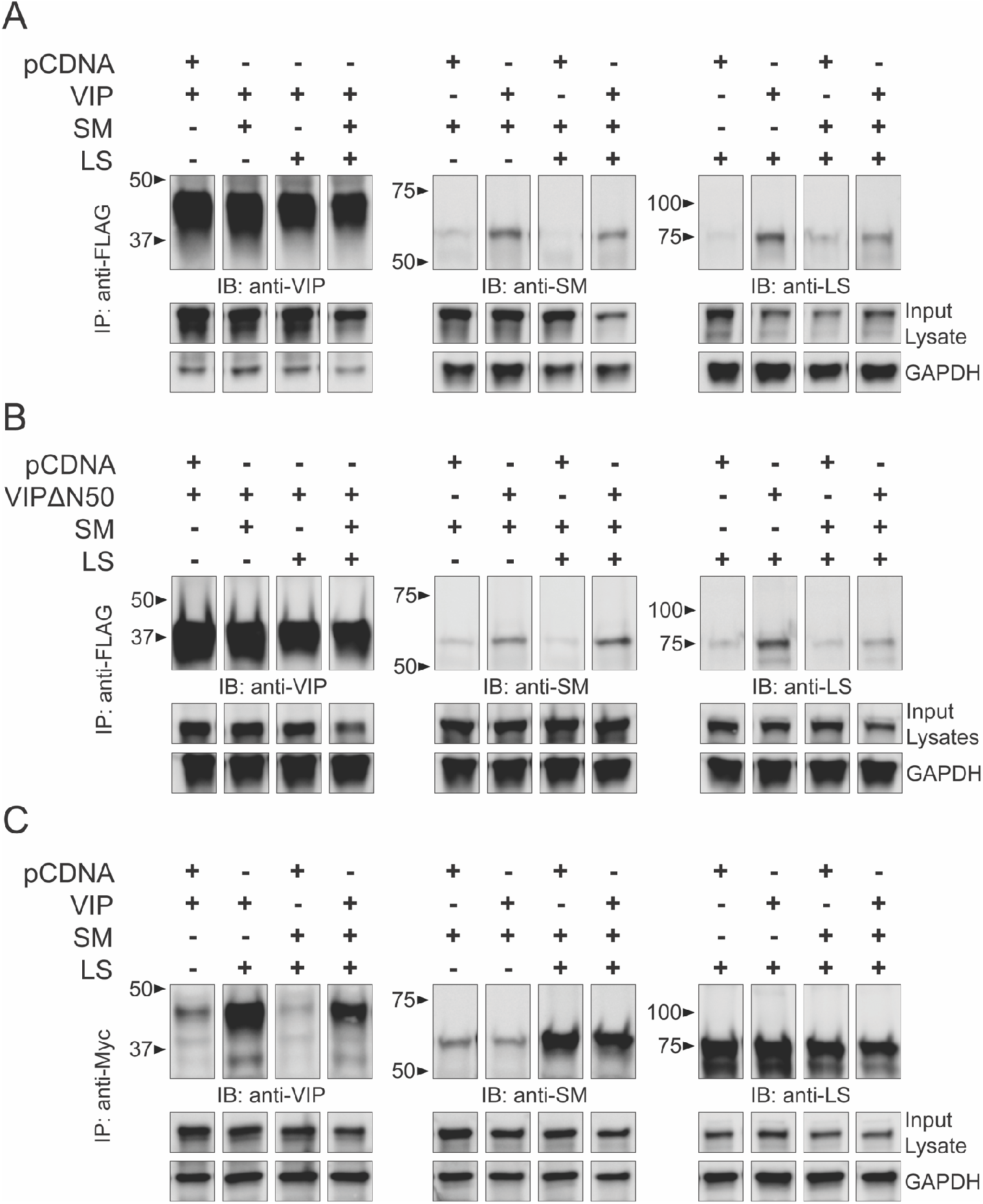
Viperin forms a ternary complex with SM and LS. **A**. HEK293T cells were transfected with empty vector (pCDNA), viperin (VIP), SM and LS as indicated. Viperin was immunoprecipitated using anti-FLAG magnetic beads and blots probed with anti-viperin, anti-SM or anti-LS polyclonal antibodies. Viperin is shown to pull down both SM and LS. **B**. The experiment described in **A**, was repeated using a viperin construct lacking the ER membrane-localizing N-terminal domain (VIPΔ50). Similar results were obtained, demonstrating that the interaction of viperin with SM and LS does not depend up ER-localization. **C.** A complementary experiment was performed in which LS was immunoprecipitated using anti-Myc magnetic beads. Consistent with the results in **A**, LS pulls down both SM and viperin.

To further validate these interactions, a second set of co-immunoprecipitation experiments was performed using LS as the bait protein and viperin and/or SM as the prey proteins. LS successfully co-precipitated both SM and viperin when each co-expressed independently with LS. LS also co-precipitated both viperin and SM when all three enzymes were co-expressed (Figure 4C). Taken together these experiments indicate that viperin binds both SM and LS, and also that SM and LS interact with each other, suggesting that the three enzymes form ternary complex formation *in vivo*. These results also indicate that the N-terminal ER-localizing domain of viperin is not required for it to bind either SM or LS.

### Viperin inhibits LS but not SM

Previous studies have shown that viperin exerts a range of effects on its interaction partners. Depending on the enzyme, viperin may inhibit or enhance catalytic activity; in other cases it may target the protein for proteolytic degradation (33, 51, 54, 55). Given the novel interaction between viperin, SM, and LS, and the important role SM and LS play in catalyzing the initial committed steps in sterol biosynthesis, we next examined whether viperin altered the enzymatic activity of either SM or LS.

SM activity assays were performed in triplicate using lysates prepared from HEK293T cells transfected with SM and the results compared with lysates prepared from cells co-transfected with viperin and/or LS. The growth medium was supplemented post transfection with BIBB 515, a potent LS inhibitor to prevent conversion of 2,3-oxidosqualene to lanosterol during the assay (62). Assays containing 0.1 mM FAD, 1.0 mM NADPH and 20 μM squalene were incubated for two hours at 37°C, after which time 2,3-oxidosqualene was extracted with ethyl acetate and quantified using LC-MS. LS assays were conducted similarly to the SM assays except that cells were not treated with BIBB 515. In this case, assays contained 50 μM 2,3-oxidosqualene as substrate and were incubated for two hours at 37°C prior to extraction and LC-MS analysis. The relative amounts of the various enzymes in the assays were quantified by western blotting, as described in the methods section.

The control SM assays exhibited an appreciable level of background activity due to endogenous SM. However, cells transfected SM exhibited a ~3-fold increase in SM activity to 7.4 ± 0.7 nmol.h^−1^.mg^−1^ of total protein, consistent with the expression of active enzyme (Figure 5A). Co-expression of viperin with SM resulted in a modest but statistically significant decrease in the level of SM expression by 29 ± 3 % (n = 3, p = 0.002; Figure S1) compared to expression of SM alone. This resulted in a corresponding decrease in the SM activity measured. But after normalizing for the changes in SM expression levels, no significant difference in the specific activity of SM was observed when it was co-expressed with either viperin or LS, or viperin and LS together (Figure 5A).

**Figure 5.**
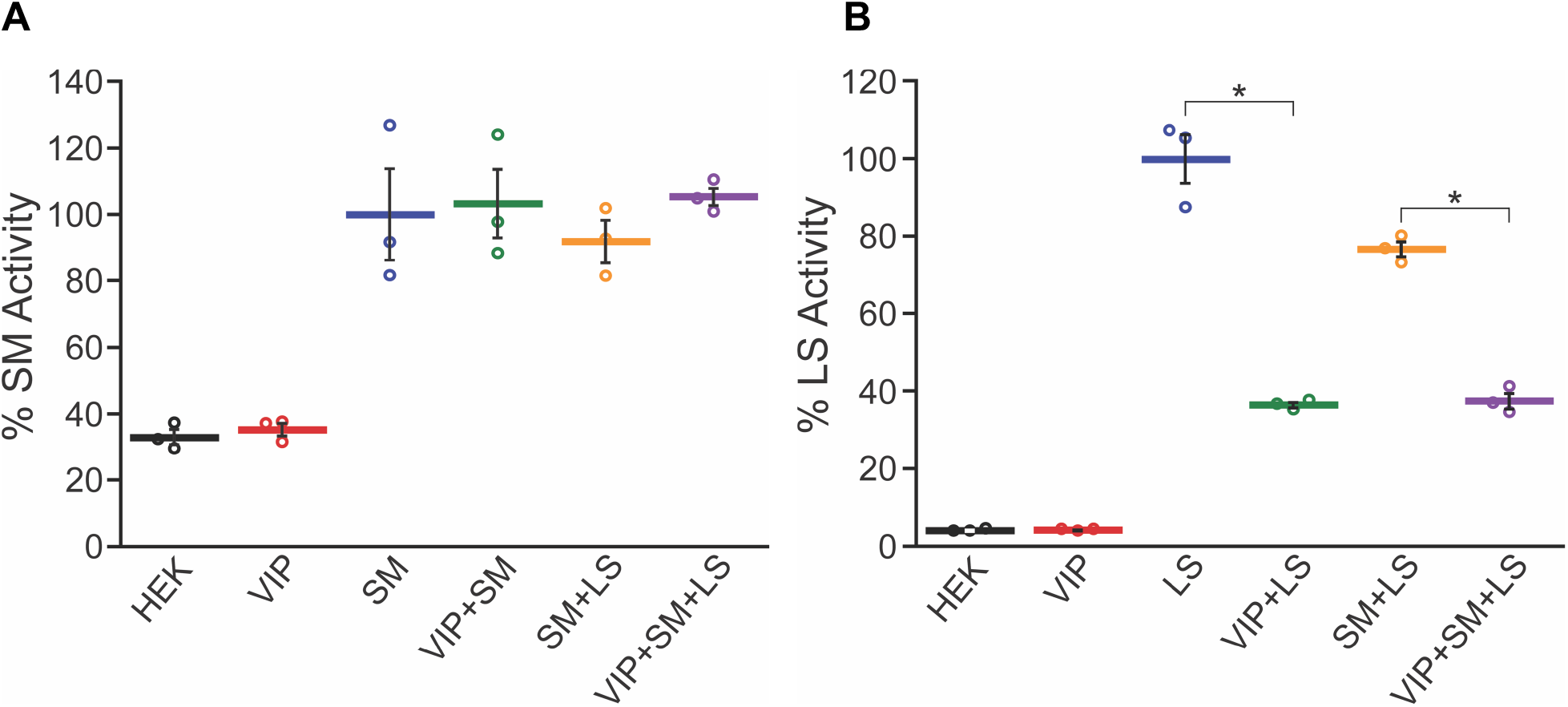
The effect of viperin expression on the enzymatic activity of SM and LS. Lysates were prepared from HEK293T cells transfected with either empty vector (HEK), viperin (VIP), SM, LS as indicated. **A.** Activity of SM. The amount of 2,3-oxidosqualene produced after two hours was determined and normalized to the amount of SM expressed in the lysate. The SM activity measured in the HEK and VIP samples arises from endogenously expressed SM. The SM-only sample is arbitrarily set as 100 %. **B.** Activity of LS. The amount of lanosterol produced after two hours was determined and normalized to the amount of LS expressed in the lysate. The LS-only sample is arbitrarily set as 100 %. Values presented are the average of three biological replicates ± SEM. * Indicates p value < 0.001.

In contrast to SM, the endogenous levels of LS activity were much lower and consequently transfection with LS resulted in a large, ~ 20-fold, increase in the LS activity of the cell lysates, with the LS activity = 6.7 ± 0.7 nmol.h^−1^.mg^−1^ of total protein (Figure 5B). In this case co-expression of viperin modestly increased the expression levels of LS by 29 ± 3 % (n = 3, p = 0.007; Figure S2) but despite this, the LS activity was substantially reduced. After normalizing for expression levels LS activity was reduced by 60 %. The inhibitory effect of viperin was independent of whether SM was co-expressed. Co-expression of SM with LS in the absence of viperin resulted in a slight decrease of ~ 20 % in LS activity that was not statistically significant, p > 0.05.

### LS inhibits ddhCTP synthesis by viperin

Our previous studies have shown that viperin’s enzymatic activity can be significantly altered by its interactions with other enzymes; in some cases viperin is activated, in others it is inhibited (43, 51). To examine whether either SM or LS potentially modulated viperin’s ddhCTP-forming activity, cell lysates were prepared under anaerobic conditions from HEK293T cells expressing either viperin alone or co-expressing SM and/or LS. Viperin activity was assayed anaerobically using 300 μM CTP and 300 μM SAM as substrates. The reaction was analyzed by quantifying the formation of 5’-deoxyadenosine (5’-dA) and the concentration of viperin in the cell extracts was determined by immunoblotting as previously described (43).

In cell lysates expressing only viperin, the observed turnover number, k_obs_ was 4.3 ± 0.5 h^−1^, which agreed well with values we previously reported (43). In lysates co-expressing SM, no significant change in viperin activity was observed, with k_obs_ = 3.7 ± 0.3 h^−1^. However, co-expression of viperin with LS resulted in a significant reduction in the specific activity of viperin, with k_obs_ = 2.2 ± 0.2 h^−1^. When all three proteins were co-expressed, viperin’s specific activity was reduced further, k_obs_ = 1.0 ± 0.2 h^−1^ (Figure 6). These data suggest that the decrease in viperin’s enzymatic activity arises from inhibition by LS that is potentiated by the interaction of viperin and LS with SM. Although these represent statistically significant decreases in viperin activity, it is unclear if the inhibitory effect of LS has any biological significance.

### Effect of LS and SM co-expression on the viperin-induced reduction in cholesterol biosynthesis

Given that viperin co-expression decreased SM levels and inhibited LS activity, we were interested to examine whether co-expression of SM or LS with viperin would reverse the decrease in cholesterol levels observed in cells expressing viperin. HEK293T cells were cultured in lipoprotein-depleted media as described above and transfected with viperin, or viperin and LS, or viperin and SM, or empty vector as a control. The cells were harvested 36 h post-transfection, as before, and total cellular cholesterol levels quantified by LC-MS. Co-expression of LS did not reverse the viperin-induced decrease in cholesterol levels, which in the viperin + LS cells were 72 ± 10 % that of the empty vector control. Similarly, co-expression of SM was also unable to reverse the viperin-induced decrease in cholesterol levels, which in the viperin + SM cells were 68 ± 2 % that of the empty vector control. Control experiments established that transfection of HEK293T cells with either SM or LS did not change cellular cholesterol levels. A further control experiment in which the viral protein, NS5A, which similarly localizes to the ER membrane, was transfected into HEK293T cells also had no effect on cellular cholesterol levels. The these experiments confirm that the observed reduction in cholesterol biosynthesis is specific to the expression of viperin.

## Discussion

Cholesterol has been shown to be essential for the viability of a number of enveloped viruses that bud from cholesterol-rich lipid rafts, including influenza A, respiratory syncytial virus (RSV) and parainfluenza viruses (18, 39, 63–66). For example, lowering cellular cholesterol levels in human alveolar epithelial (A549) cells by treatment with either gemfibrozil and/or lovastatin significantly impairs the budding of human parainfluenza virus, thereby reducing viral titers by up to 98 % (67). In the case of influenza A and RSV, viral titers were reduced less dramatically, by ~ 10-fold, but the resulting viral particles, which contained reduced amounts of cholesterol, were both less stable and less infectious (67). These observations demonstrate the potential importance of down-regulating cholesterol biosynthesis as an antiviral response to enveloped viruses. Our results show that viperin expression lowers cellular cholesterol levels by between 20 – 30 %, depending upon the experimental conditions, which, based on these prior studies, would substantially impair viral budding and viability.

The initial indication that viperin may down-regulate cholesterol biosynthesis came from studies in which viperin was found to retard influenza A virus particles from budding from infected cells, although changes in cholesterol levels were not reported (18, 39). Using a yeast 2 hybrid screen, viperin was identified as interacting with farnesyl pyrophosphate synthase (FPPS), which catalyzes an early step in the cholesterol biosynthetic pathway.(18) However, more recent studies from our laboratory failed to identify a physical interaction between FPPS and viperin. Rather, it appears that viperin may indirectly down-regulate FPPS expression, likely by increasing its rate of proteolytic degradation (55). Consistent with these earlier results, FPPS was not identified in the proteomic screen described here. Nevertheless, viperin-induced down-regulation of FPPS may contribute to overall decrease in cholesterol biosynthesis we observed.

Multiple studies have provided evidence for viperin’s interaction with a remarkably wide range of proteins (19, 21, 68). Many of the protein partners for viperin have been inferred by following changes in cellular physiology upon infection with various viruses. Here, we sought to achieve a broader picture of the proteins that may interact with viperin using a proteomics approach. The identification of several proteins that have previously been documented to interact with viperin, such as HADHA and HADHB (fatty acid β-oxidation) (37), CIAO1 and MMS19 (iron-sulfur cluster installation) (69) serves to validate the screening approach. However, we note that protein partners that are in low cellular abundance may have been missed by the screen, as would proteins that are only expressed upon viral infection. Also, the location of viperin at the ER-membrane may bias the analysis towards enrichment of other ER-resident proteins. In some respects, our results reflect the consensus in the literature that viperin appears to be fairly promiscuous in its interactions with other proteins (36, 40, 49), as evidenced by the wide variety of proteins represented in the list of potential interaction partners. However, in addition to cholesterol biosynthetic enzymes, our analyses using DAVID also identified that a significant number of proteins involved in membrane lipid metabolism interact with viperin (Table S1). This observation suggests that, beyond lowering cholesterol levels, viperin may alter lipid metabolism and hence the lipid composition of cell membranes more extensively in response to viral infections.

The two enzymes that viperin appears to interact with most strongly, based on their enrichment factors, are LS and SM. SM catalyzes the committed step in sterol biosynthesis and is highly regulated by both transcriptional and post-translational mechanisms (10–12). Indeed most, if not all, cholesterol biosynthetic enzymes appear to be regulated at the transcriptional level in response to sterol levels through the action of SREBPs (3, 4, 9, 70). Although the regulation of LS is less well studied, the fact that SM and LS form a complex with other cholesterol biosynthetic enzymes (59) suggests that the effects of regulating SM may be propagated to other enzymes in the complex. The cholesterol biosynthetic enzymes reside in the ER membrane and are therefore likely to be in close proximity to viperin, which is also localized to the ER membrane. It therefore seems reasonable that these enzymes could be subject to regulation by viperin.

We consistently observed that co-expression of viperin reduced the levels of SM accumulating in cells by ~ 30%. Although this is not a large change in protein levels, it does suggest that viperin may reduce SM levels by increasing its rate of proteasomal degradation. In support of this idea, the ubiquitin-dependent degradation of SM is known to be regulated in response to cellular cholesterol levels through the action of the E3 ubiquitin ligase MARCH6 (11) and it is generally considered that viperin exerts some of its antiviral effects by increasing the rate of proteasomal degradation of its target proteins (15, 68). Although the evidence for viperin’s interaction with proteasomal degradation machinery is indirect, viperin is known to activate TRAF6, which is an E3 ligase involved in the K63-linked polyubiquitination of proteins involved in immune signaling (49, 51).

Although we observed that viperin co-expression reduced the enzymatic activity of LS in cell lysates and the expression levels of SM, co-expression of either LS or SM with viperin did not reverse the viperin-induced decrease in cellular cholesterol levels. This observation was somewhat surprising, given the strong interactions between these enzymes and viperin and their important roles in cholesterol biosynthesis. However, it is possible that under the conditions of these experiments, in which LS, SM and viperin were expressed at artificially high levels, that there was still sufficient viperin present to titrate out the additional LS and SM. It is also possible that viperin down-regulates cholesterol biosynthesis by restricting the flux through other points in the pathway, for example by reducing FPPS levels, or through a more circuitous route not captured in these studies. Further experiments will be needed to better establish how viperin down-regulates this important biosynthetic pathway.

## Conclusions

We observed that transient expression of viperin results in ay 20 – 30 % decrease in cellular cholesterol biosynthesis, which is sufficient to explain the previously observation that viperin retards virus budding from the cell membrane. Consistent with this observation, the interactome of viperin includes a number of enzymes involved in the later stages of cholesterol biosynthesis. Of these, SM and LS were the most highly enriched proteins identified and their interactions with viperin were validated by co-immunoprecipitation. Co-expression of viperin reduced the cellular levels of SM by ~ 30 % and inhibited the enzymatic activity of LS by over 60 %. However, over-expression of either SM or LS in HEK293T cells failed to reverse the effects of viperin expression on cellular cholesterol levels. This observation suggests that viperin’s down-regulation of cholesterol biosynthesis is a complex phenomenon and not simply an effect of its interactions with these two biosynthetic enzymes.

## Experimental procedures

### Cell Lines

HEK 293T cells were obtained from ATCC.

### Plasmids

The expression constructs for viperin and viperin-ΔN50, lacking the N-terminal 50 residues, in pcDNA3.1(+) were as described previously.(43) The genes encoding SM and LS were synthesized and cloned into the pcDNA3.1(+) vector commercially (GenScript). SM was inserted between the BamHI and XbaI restriction sites and included a C-terminal V5 tag. LS was inserted between the HindIII and BamHI restriction sites and included a C-terminal Myc tag. A Kozak consensus sequence (5’-GCCACC-3’) was included upstream of each protein to allow for protein expression in mammalian (HEK293T) cells.

### Antibodies

Rabbit polyclonal viperin (11833-1-AP), mouse monoclonal viperin (MABF106), goat anti-rabbit Ig secondary (170-6515), and goat anti-mouse Ig secondary (626520) antibodies were used as described previously.(43) Rabbit polyclonal squalene monooxygenase (12544-1-AP), rabbit polyclonal lanosterol synthase (13715-1-AP), and rabbit polyclonal GAPDH (10494-1-AP) antibodies were purchased from Protein Tech. Mouse monoclonal GAPDH (6C5) antibody (CB1001) was purchased from EMD Millipore.

### Reagents

For cell culture, Dubelco’s modified eagle medium (DMEM), Opti-MEM reduced serum medium, 0.05% Trypsin-EDTA and phosphate buffered saline were obtained from Gibco (ThermoFisher Scientific); lipoprotein deficient serum from fetal calf was purchased from Millipore Sigma. For transfection of DNA into HEK293T cells, transfection grade linear polyethyleneimine HCl was purchased from Polysciences, Inc and Fugene transfection agent was purchased from Promega. Phenylmethylsulfonyl fluoride (PMSF) and Roche complete EDTA-free protease inhibitor cocktail tablets were purchased from ThermoFisher Scientific and Millipore Sigma, respectively. For co-immunoprecipitation, anti-FLAG M2 magnetic beads and Pierce anti-c-Myc magnetic beads were purchased from Millipore Sigma and ThermoFisher Scientific, respectively. S-(5′-Adenosyl)-L-methionine p-toluenesulfonate salt was purchased from Millipore Sigma, and CTP disodium salt hydrate was purchased from Acros Organics. For the SM and LS assays, NADPH tetrasodium salt, flavin adenosine dinucleotide disodium salt hydrate, 2,3-oxidosqualene, lanosterol, and 2,6-Di-tert-butyl-4-methylphenol (BHT) were purchased from Millipore sigma. Squalene was purchased from ThermoFisher Scientific and BIBB 515 was purchased from Cayman Chemical.

### Cell Culture, Transfection, Harvesting

HEK293T cells were cultured, transfected, and harvested as described previously.(43) Briefly, HEK293T cells were grown to 50-60% confluency after which transfection with the desired construct(s) (in pcDNA3.1(+)) was carried out using a polyethyleneimine (PEI) transfection agent. Transfections (and co-transfections) were performed with 20 μg each plasmid, using a 1:2 ratio of plasmid to PEI. DNA and PEI were mixed and incubated for 10 minutes at room temperature then added to HEK293T cells (at the desired confluency) in a 10 cm culture plate. Typically cells were allowed to grow for a further 24 - 48 hours before harvesting by gentle centrifugation and stored at −80°C. For cells used in squalene monooxygenase assays, the lanosterol synthase–specific inhibitor, BIBB 515, 1 μM was added 24 hours post transfection to prevent further conversion of 2,3-oxidosqualene to lanosterol by endogenous LS(62).

The cells used for cholesterol analyses were first gradually acclimated to increasing Opti-MEM serum-free media in four passages at 10, 25, 50, and 75% Opti-MEM media, to allow for normal growth with minimal serum. Then cells were passed into 90% Opti-MEM medium and 10% medium comprising DMEM supplemented with lipoprotein depleted fetal calf serum; grown to 50-60% confluency and transfected with either an empty pCDNA3.1 (+) vector as a control or viperin in pCDNA3.1 (+) as described earlier. Cells were grown for 36 hours post-transfection, harvested, and stored at −80°C.

### Proteomic screening

HEK293T cells over-expressing either 3x-FLAG-viperin or 3x-FLAG-tagged empty vectors (3x-FLAG-pcDNA3.1) as a control were harvested in PBS from 10 cm plates. Cells were incubated on ice for 20 minutes with 0.5 mL lysis buffer (20mM Tris pH 7.5, 500mM NaCl, 0.1% Tween-20, 1 mM phenylmethylsulfonyl fluoride, Complete Protease Inhibitor Cocktail) and lysed using a Fisher Scientific handheld sonicator at 10% amplitude (25 pulses 1 second on, 5 seconds off). Cell lysates were cleared by centrifugation at 14,000 RPM for 30 minutes at 4°C and the pellets were discarded. Anti-FLAG M2 Magnetic Beads (Sigma Aldrich, M8823) were washed and pre-equilibrated with ice-cold 20mM Tris pH 7.5, 500mM NaCl, 0.1% Tween-20. Total protein concentration of the supernatant was measured by Protein DC Assay (Bio-Rad) and subjected to immunoprecipitation using anti-Flag M2 magnetic beads. Supernatant (total protein concentration ~ 7 μg/μL) was incubated with the pre-equilibrated anti-FLAG magnetic beads at 50:1 (w/v) ratio for two hours at 4°C. Beads were washed three times with 20x bead volume of TBS using a magnetic tube rack (MagRack 6, GE Life Sciences).

The beads were re-suspended in 50 μL of 0.1M ammonium bicarbonate buffer (pH~8); cysteine residues were reduced by adding 50 μL of 10 mM DTT and incubating at 45° C for 30 min. Samples were cooled to room temperature and cysteine alkylated by incubation with 65 mM 2-chloroacetamide for 30 min. at room temperature in the dark. Overnight digestion with 1 μg sequencing grade, modified trypsin was carried out at 37° C with constant shaking. Digestion was stopped by acidification and peptides were desalted using SepPak C18 cartridges (Waters). Samples were dried and the resulting peptides were re-dissolved in 8 μL of 0.1% formic acid/ 2% acetonitrile solution. 2 μL of the peptide solution were chromatographed on a nano-capillary reverse phase column (Acclaim PepMap C18, 2 micron, 50 cm, Thermo Scientific) using a 0.1% formic acid/ 2% acetonitrile (Buffer A) and 0.1% formic acid/95% acetonitrile (Buffer B) gradient at 300 nL/min over a period of 180 minutes (2-22% buffer B in 110 minutes, 22-40% in 25 minutes, 40-90% in 5 minutes followed by holding at 90% buffer B for 5 minutes and re-quilibration with Buffer A for 25 minutes). Eluent was directly introduced into Orbitrap Fusion Tribrid mass spectrometer (Thermo Scientific, San Jose CA) using an EasySpray source. MS1 scans were acquired at 120K resolution (AGC target=1×106; max IT=50 ms). Data-dependent collision induced dissociation MS/MS spectra were acquired using Top speed method (3 seconds) following each MS1 scan (NCE ~32%; AGC target 1×105; max IT 45 ms).

Proteins were identified by comparing the MS/MS data against *Homo sapiens* protein database (UniProt; 42054 entries, 2016-11-30) using Proteome Discoverer (v2.1, Thermo Scientific). Search parameters included MS1 mass tolerance of 10 ppm and fragment tolerance of 0.2 Da; two missed cleavages were allowed; carbamidimethylation of cysteine was considered a fixed modification and oxidation of methionine, deamidation of asparagine and glutamine, phosphorylation of serine, threonine and tyrosine, ubiquitination of lysine (diglycine signature) were considered as potential modifications. False discovery rate (FDR) was determined using Percolator and proteins/peptides with an FDR of ≤1% were retained for further analysis.

### Immunoblotting

Immunoblotting of HEK293T cell lysates was performed as previously described.(51, 55) Primary antibodies were used at the following dilutions: rabbit polyclonal and mouse monoclonal anti-viperin diluted 1:3000, rabbit polyclonal anti-SM diluted 1:2000, rabbit polyclonal anti-LS diluted 1:1000, rabbit polyclonal and mouse monoclonal anti-GAPDH diluted 1:5000. Both goat anti-rabbit and goat anti-mouse Ig secondary antibodies were diluted at 1:5000. All dilutions were done using 5% w/v skim milk dissolved in 20mM Tris pH 7.5, 500mM NaCl, 0.1% Tween-20. All quantitative protein measurements presented represent the average of at least three independent biological replicates in coordination with GAPDH controls.

### Co-immunoprecipitation Assays

Immunoprecipitation experiments were performed essentially as described previously(43, 51). When using viperin as the bait protein, Anti-FLAG M2 Magnetic Beads (Sigma Aldrich, M8823) were pre-equilibrated with 20 mM Tris/Cl pH 7.5, 500 mM NaCl, 0.1% Tween-20 (wash buffer). 50 μL of 50% slurry (per culture) was added to cleared cell lysate in a 1.5 mL Eppendorf tube and incubated by end-to-end mixing for two hours at 4°C. The flow through was removed by placing the tube in a magnetic tube rack for one minute, waiting for all beads to migrate to the side of the tube, and removing the residual solution. The beads were washed with wash buffer three times for five minutes each by end-to-end mixing at 4°C. Protein complexes were eluted by adding 40 μL of 4x sample buffer supplemented with 2 mM 2-mercaptoethanol to each sample and subsequent incubation at 95°C for five minutes. The samples were centrifuged to remove the magnetic beads and analyzed by immunoblotting. When using LS as the bait protein, the same protocol was used except that 40 μL of 50% slurry (per culture) of pre-equilibrated and chilled Pierce Anti-c-Myc Magnetic Beads (ThermoFisher Scientific) were substituted for anti-FLAG beads.

### Viperin activity assays

Viperin activity assays were performed as previously described.(43) All assays were carried out using at least three biological replicates in an anaerobic chamber (COY) with O_2_ content < 50 ppm. Briefly, HEK293T cells containing viperin and/or SM and/or LS were harvested from one 10 cm diameter tissue culture plate for each assay and re-suspended in 0.5 mL anoxic lysis buffer (above). The cell suspension was lysed by sonication and centrifuged for 15 minutes at 14,000 r.p.m. in an Eppendorf centrifuge at 4°C. The supernatant was used for the activity assay. 5 mM dithiothreitol (DTT), 5 mM sodium dithionite, and 300 μM CTP were added to the cleared lysate and incubated at room temperature for 30 min. The reaction was initiated by the addition of 200 μM S-adenosyl-L-methionine (SAM) and allowed proceed for 1 h at room temperature. The reaction was quenched by incubation at 95°C for 5 min. After chilling the solution to 4°C, it was centrifuged for 30 minutes at 14,000 r.p.m. and the supernatant was removed. The organic component of the solution was extracted with acetonitrile and analyzed in triplicate by UPLC-MS/MS as previously described.(71)

### Squalene monooxygenase activity assays from HEK293T cell lysate

HEK293T cells overexpressing different combinations of viperin, SM, and LS (which had been incubated post-transfection with 1 μM BIBB 515) were harvested from a 10 cm diameter tissue culture plate; the cells from one plate were sufficient for one assay. HEK293T cells transfected with an empty pcDNA3.1(+) vector were used as a negative control. SM activity was assayed by measuring the conversion of squalene to 2,3-oxidosqualene. Cells were re-suspended lysis buffer (TBS, 0.1% Tween 20, complete protease inhibitor cocktail) and lysed as described above. All assays were performed in lightproof tubes and completed as previously described(72) with minor modification. Briefly, assays contained 200 μL cleared HEK cell lysate from cells expressing the protein(s) of interest; 0.1 mM FAD, 1 mM NADPH, and were initiated by the addition of squalene (500 μM stock, all solutions prepared in ethanol), final concentration 20 μM in a final reaction volume of 250 μL. Assays were incubated at 37°C for 2 h and quenched with 1 mL ethyl acetate supplemented with 0.2 mg/mL BHT purged with N_2_ followed by vortexing at room temperature for 30 min. The organic layer was isolated by centrifugation for 10 minutes at 14,000 rpm. 300 μL of the organic layer was removed and evaporated to complete dryness under a nitrogen atmosphere. The resulting film was reconstituted with 50 μL acetonitrile supplemented with 0.2 mg/mL BHT.

LC-MS analysis was performed as previously described(72) with minor differences. Briefly, 2 μL of sample was injected into and Agilent 1290 Infinity Series HPLC equipped with a Waters Acquity UPLC BEH C18 column. (1.7 um, 2.1×50 mm) in line with an Agilent Q-TOF Mass Spectrometer (model G6520B). The HPLC separation was done on a Waters Acquity UPLC BEH C18 column. (1.7 um, 2.1×50 mm). Mobile phase A comprised water with 0.1% formic acid while mobile phase B comprised 20% isopropanol, 80% acetonitrile, and 0.1% formic acid. Analytes were eluted isocratically at 0.6 mL/min using 2% mobile phase A and 98% mobile phase B over 3.75 min. Q-TOF spectra were acquired using an APCI ion source in positive ion mode. The mass spectrometer was scanned from m/z 50 to m/z 1000 with one spectrum per second and 2,3-oxidosqualene was detected by monitoring a mass transition pair of m/z 427.38 to 409.38. The LC-MS chromatograms were integrated and resulting peaks were analyzed using Mass Hunter software. Measurements represent the average of at least three biological replicates, with three technical replicates of each.

### Lanosterol synthase activity assays from HEK293T cell lysate

LS activity assays were carried out in the same manner as the SM assays except that cells were not treated with BIBB 515. HEK293T cells transfected with an empty pcDNA3.1(+) vector were used as a negative control. For each assay cells were harvested from a single 10 cm diameter tissue culture plate and cleared lysates prepared. LS activity was measured by formation of lanosterol from 2,3-oxidosqualene. All reactions were carried out in lightproof tubes and contained 200 μL cleared HEK cell lysate and were initiated by the addition of 50 μM 2,3-oxidosqualene, final concentration (500 μM stock, all solutions prepared in ethanol), in a final reaction volume of 250 μL. Assays were incubated in a 37°C for 2 h and then worked up as described above except the reactions were quenched with chloroform. LC-MS analysis was performed as described above, but the ion analyzed was that of the lanosterol radical without hydroxide at m/z 409.38. Measurements represent the average of at least three biological replicates, with three technical replicates of each.

### Cholesterol Analysis of HEK293T cells

HEK293T cell pellets were re-suspended in 500 μL TBS supplemented with 0.1% Tween-20 (TBST) and lysed by sonication. Cholesterol was extracted from 50 μL of the resulting lysates with 1.2 mL ethyl acetate supplemented with 0.2 mg/mL BHT purged with N_2_ and vortexing at room temperature for 30 min. The organic component was isolated by centrifugation for 10 minutes at 14,000 rpm. 500 μL of the organic layer was removed and evaporated to dryness under a nitrogen atmosphere. The resulting film was redissolved in 500 μL acetonitrile containing 0.2 mg/mL BHT. LC-MS analysis was performed as described above, except the ion analyzed was that of the cholesterol radical without hydroxide at m/z 369.25. Cholesterol content was determined with respect to a standard curve constructed from pure cholesterol samples. Cholesterol concentrations were normalized with respect to the total protein concentration of the cell determined by BCA assay. Measurements generally represent the average of at least three biological replicates, with three technical replicates of each.

## Acknowledgements

This work was supported in part by NIH grants GM 093088 to E.N.G.M. and DK 046960 to R.T.K.

## Author Contributions

T.J.G., S.G., A.M.P., K.S., J.W., V.B. and Y.S. performed the experiments. T.J.G., S.G., A.M.P., V.B., A.I.N. and Y.S. analyzed the results. T.J.G., S.G., A.I.N., R.T.K. and E.N.G.M. designed the experiments. T.J.G., S.G. and E.N.G.M. wrote the paper.

